# *LsTT2* encoding R2R3-MYB transcription factor is responsible for a shift from black to white in lettuce seed

**DOI:** 10.1101/2023.09.17.558082

**Authors:** Kousuke Seki, Kenji Komatsu, Kanami Yamaguchi, Yoshinori Murai, Keiji Nishida, Ryohei Koyama, Yuichi Uno

## Abstract

Prickly lettuce (*Lactuca serriola*), which is considered the wild ancestor of lettuce, has black seeds, whereas the major seed color of domesticated lettuce is black or white. The successfully-selected white seed trait is a key domestication trait for lettuce cultivation and breeding; however, the mechanism underlying the shift from black to white seeds remains to be clarified. We aimed to identify the gene/s responsible for white seed trait in lettuce. Genetic mapping of a candidate gene was performed with double-digest RAD sequencing using an F_2_ population derived from a cross between ‘ShinanoPower’ (white seed) and ‘Escort’ (black seed). The white seed trait was controlled by a single recessive locus (48.055–50.197 Mbp) in linkage group 7. Narrowing down using five PCR-based markers and 84 cultivars, eight candidate genes were mapped in the locus. Only the *LG7_v8_49*.*251Mbp_HinfI* marker, which employs a single nucleotide mutation in the stop codon of *Lsat_1_v5_gn_7_35020*.*1* was completely linked to the seed color phenotype. In addition, the sequences of the coding region for candidate genes except for *Lsat_1_v5_gn_7_35020*.*1* were identical in the resequence analysis of ‘ShinanoPower’ (white seed) and ‘Escort’ (black seed). Therefore, we proposed *Lsat_1_v5_gn_7_35020*.*1*, a gene located in the locus, as the candidate gene and designated it as *LsTT2*, an ortholog encoding the R2R3 MYB transcription factor in *Arabidopsis*. When we validated the role of *LsTT2* in seed color through genome editing, *LsTT2* knockout mutants harboring an early termination codon showed a change in seed color from black to white. White seeds accumulated less proanthocyanidins than black seeds, which was similar to the phenotype observed in *Arabidopsis TRANSPARENT TESTA 2* (*TT2*) mutants. Therefore, *LsTT2* was the allele responsible for the shift in seed color from black to white. The development of a robust marker for marker-assisted selection and identification of the gene responsible for white seeds has implications for future breeding technology and physiological analysis.

## 1 Introduction

In lettuce (*Lactuca sativa* L.), the major seed color are black and white. Black-colored seed is the wild-type trait observed in *Lactuca serriola* L., a wild lettuce species distributed worldwide. The white-colored seed trait was possibly discovered during the process of domestication. We infer that individual plants with extremely reduced pigments in the seed pericarp, where the pigments are localized in lettuce, was accidentally discovered (Thompson, 1942). Black seeds are difficult to identify on the soil but white seeds could be easily identified on the soil (Fig. 1a and 1b). Lettuce seeds are small; therefore, the easy visibility that a white seed offers is an advantage in seeding and harvesting. White seeds are desirable for agricultural production. Lettuce seeds are sown near the soil surface as a standard practice (Woolley and Stoller, 1978), because lettuce seed germination is promoted by light radiation (Borthwick et al., 1952). Light transmission at 660 and 730 nm is involved in the germination of black and white seeds, respectively. Transmission spectra of black seeds show transmission of less than 20% of the incident light, whereas transmission spectra of white seeds indicate a transmission of more than 50% of the incident light (Widell and Vogelmann, 1988). In addition, white seeds were more sensitive to temperature than black seeds, allowing white seeds to germinate in darkness (Borthwick et al., 1952). Therefore, white seeds are believed to be more advantageous and exhibit better performance and germination than black seeds. The white seed trait is common and is present in well-known cultivars, such as cv ‘New York’. Several white seed varieties are available in the database; therefore, we believe that the varieties were bred through artificial selection (Table S1). Though the white seed trait is recessive (Thompson, 1942; Ryder, 1999; Wang et al., 2016), the data of seed lists of the Centre for Genetic Resources, the Netherlands (CGN: https://www.wur.nl/en/Research-Results/Statutory-research-tasks/Centre-for-Genetic-Resources-the-Netherlands-1.htm) and the Germplasm Resources Information Network (GRIN: https://www.ars-grin.gov/) show that the number of white seed cultivars is significantly higher than that of black seed cultivars (Table 1). This fact implies that lettuce breeders around the world intentionally have introduced the trait of white seed into new breeding cultivars. The white seed is an important agricultural trait for lettuce breeders; however, the molecular mechanism underlying the shift from black to white remains incompletely understood (Thompson, 1942; Waycott et al., 1999; Kwon et al., 2013). White seed trait is controlled by a recessive single gene (Waycott et al., 1999) located on LG7 (Kwon et al., 2013). We applied the ddRAD-seq method to analyze the genetic details of the white seed. Further mapping was performed using 84 cultivars to narrow down the gene, and the only candidate gene identified was validated for involvement in seed color using a knockout mutant through genome editing.

**Table 1.** Number of black seed and white seed cultivars of lettuce germplasm.

| Plant Resource | Total | Black seed | White seed |
| --- | --- | --- | --- |
| CGN | 1,453 | 512 | 941 |
| GRIN | 2,441 | 1,127 | 1,314 |
| CGN + GRIN | 3,894 | 1,639 | 2,255 |

**Fig. 1.**
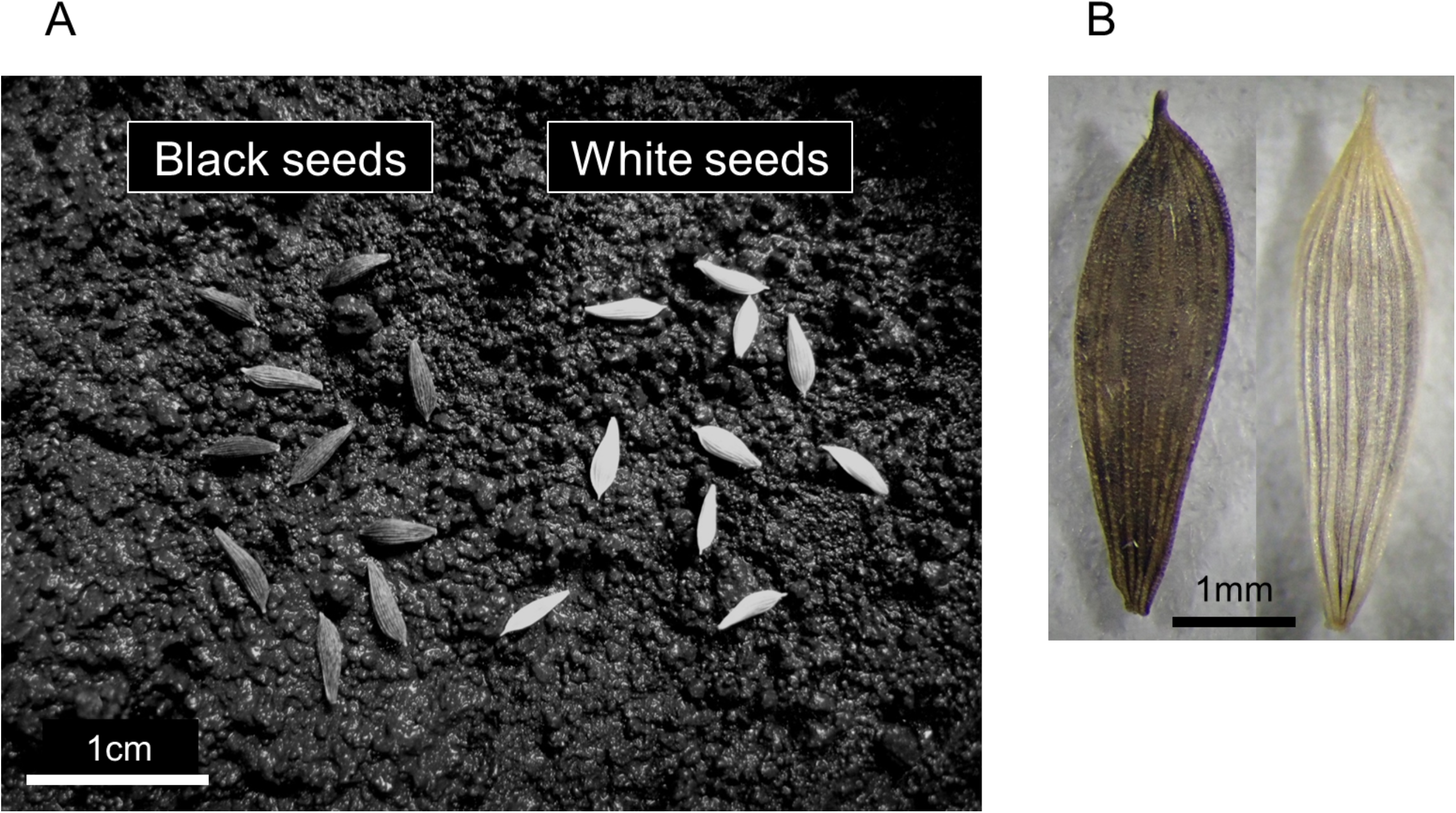
Comparison of black and white seeds of lettuce. (A) Black and white seeds shown on the soil surface. (B) Black and white seeds shown on a background of intermediate color.

## 2 Materials and methods

### 2.1 Plant material

The crisphead-type lettuce cultivar, ‘ShinanoPower’ was bred by the Nagano Vegetable and Ornamental Crops Experiment Station, and ‘Escort’ was bred by Takii & Co., Ltd. ‘ShinanoPower’ and ‘Escort’ were crossed to produce an F_1_, which was selfed. Approximately 96 F_2_ individuals were investigated for seed color in the greenhouse. The oilseed-type lettuce cultivar ‘Oilseed’ derived from upper Egypt was introduced from CGN; the original strain number is ‘CGN04769’.

### 2.2 Double-digest RAD sequencing (ddRAD-Seq) and resequencing

Genomic DNA was extracted from the leaves using a NucleoSpin Plant II Extract Kit (Machery-Nagel, Duren, Germany). ddRAD-seq and resequencing were performed as described by Seki et al. (2020) and Seki (2021). The ddRAD-seq libraries were sequenced using the Illumina Hiseq4000 platform. Paired-end sequencing reads (100 bp x2) were analyzed for ddRAD-seq tag extraction, counting, and linkage map construction using RAD-R scripts (Seki, 2021). The linkage map was graphically visualized using MapChart (Voorrips, 2002). Resequencing libraries were sequenced on the HiSeqX platform. DNA samples from the two parental lines were used to construct paired-end sequencing libraries (150 bp x2) and were subjected to whole-genome sequencing. Raw sequence data (fastq) for the present RAD-seq and resequence analysis are available in the DNA Data Bank of Japan (DDBJ) Sequence Read Archive (http://ddbj.nig.ac.jp/dra/index_e.html) under accession number DRA010289.

### 2.3 Resequence analysis of parent genomes and sequencing of a candidate locus

Resequencing reads from the two parental cultivars were mapped onto the reference genome *L. sativa* cv Salinas V8 using the BWA software (Li and Durbin, 2009). A detailed script is provided in Resequence_mapping_script_BWA_mem.txt (https://github.com/KousukeSEKI/RAD-seq_scripts). The sorted BAM files were visualized using the IGV software (Robinson et al., 2011).

### 2.4 Genotyping using publicly available genome resequencing data

The publicly available *L. sativa* genome sequencing data were obtained from the NCBI Sequence Read Archive (SRA). FASTQ files were imported into the CLC Genomics Workbench (QIAGEN, USA) for subsequent analysis. The trim sequence tool in the suite was used to filter out low-quality bases (<Q30), and only reads that showed a quality score of ≥30 were retained. Filtered sequence reads were mapped onto the *L. sativa* v8.0 genome (https://phytozome.jgi.doe.gov/pz/portal.html#!info?alias=Org_Lsativa_er) using the Map Reads Reference tool, and Local Realignment was performed using the Local Realignment tool. Based on the mapping results, the stop codon polymorphism in the *LsTT2* was identified.

### 2.5 Designing PCR-based markers and their amplification

Polymorphisms near the locus at 48–50 Mbp in LG7, including insertions, deletions, and single-nucleotide polymorphisms (SNPs), were evaluated as potential markers. The primer names were formatted as (linkage group) _ (genome version) _ (genome position) _ (restriction enzyme, in case of CAPS). Primers for locus amplification were designed using Primer3 (http://bioinfo.ut.ee/primer3-0.4.0/), and KOD FX (TOYOBO, Japan) was used for amplification. PCR was performed using 0.5 μL of DNA template, 0.4 μL of each primer (50 μM), 2 μL of dNTP (2 mM), 5 μL of 2× PCR Buffer, 0.2 μL of KOD FX (1 U/μL), and distilled water (dH_2_ O) to a final reaction volume of 10 μL. The PCR conditions were as follows: 94°C for 5 min, 30 cycles of 94°C for 30 s and 62°C for 30 s, followed by one cycle at 72°C for 4 min. After amplification, electrophoresis was performed using 9 μL of the PCR products on a 2% agarose gel (Takara Bio, Japan) at 100 V. In case of CAPS, the PCR products were digested at 37°C for 1 h in 20 µl total volume with 5−15 units of the appropriate restriction enzyme before electrophoresis.

### 2.6 Genome editing

The 20-nt gRNAs specific for the target gene, *LsTT2* were designed for the first exon (Fig. S1 and Table S2). Primers with the *Bbs*I restriction site were annealed and ligated into the entry vector, 1480_MluI-1433_pUC19_AtU6oligo (Shimatani et al., 2017). The core partial fragments were excised from the entry vector using *I-Sce*I and subcloned into the T-DNA region of the destination vector, 1432_pZD_OsU3gYSA_HolgerCas9_NPTII (Shimatani et al., 2017). All vectors were constructed through standard cloning and verified through sequencing. Lettuce ‘Oil seed’ was transformed with Agrobacterium harboring the destination vector. Regenerated plants were selected using kanamycin, and genome integration of T-DNA was confirmed through PCR using NPTII. To determine the genome editing ability of *LsTT2*, the target sequence was amplified from gDNA through PCR and cloned into the standard sequencing vector, pSKII^-^. Eight clones of each T_0_ strain were used in this study.

### 2.7 Analysis of the pigment in seed

White and black seeds were frozen in liquid nitrogen and powdered. The samples were stored at -40°C until use. The proanthocyanidin content of each seed was compared using vanillin–sulfuric acid assay modified from Sugawara et al.

(https://www.naro.affrc.go.jp/project/results/laboratory/karc/2004/konarc04-22.html; accessed February 10, 2020). The freeze-dried powder (100 mg) of each seed was extracted using 1 mL of methanol with shaking. The extracts were mixed with 2 mL 1% (w/v) vanillin/methanol, and 2 mL 25% (v/v) sulfuric acid/methanol was added to the solution and shaken at 30°C for 15 min. An additional 1 mL of methanol was added to the solution. After centrifugation (3000 rpm/min, 10 min), the absorbance of the supernatant was measured at 500 nm using a Shimadzu UV-2600 UV-Vis spectrometer (Shimadzu Corporation, Japan). To characterize the composition of phenolic compounds, including anthocyanins and other flavonoids, seed powder (100 mg) was extracted with formic acid/H_2_ O/methanol (5:10:85, v/v/v). The extracts were filtered with a GL Chromatodisk 13N (0.45 μm pore size, GL Sciences, Inc., Japan) and analyzed using a Shimadzu Prominence HPLC system with a SunShell C18 column [2.6 μm particle material, I.D. 4.6 × 100 mm (ChromaNik Technologies Inc., Japan)] at a flow rate of 0.4 mL/min, detection wavelength of 190–700 nm, and eluent: phosphoric acid/acetonitrile/acetic acid/H_2_ O of 3:6:8:83 (v/v/v/v). The injection volume was 1 μL.

## 3 Results

### 3.1 Inheritance of seed color

An F_2_ population was derived from an initial cross between ‘ShinanoPower’ (white seed) and ‘Escort’ (black seed) to elucidate the inheritance of the white seed trait. The F_1_ plants produced black seeds. Among the 96 F_2_ individuals, 74 possessed black seeds and 22 possessed white seeds (Table 2). These results suggest that the white-seed trait is determined by a single recessive locus.

**Table 2.** χ2 test for the seed color of F_2_ populations.

| Population | Total | Black seed | White seed | Segregation ratio | $\chi^2$<br>(3:1) |
| --- | --- | --- | --- | --- | --- |
| Escort | 10 | 10 | - | - | - |
| ShinanoPower | 10 | - | 10 | - | - |
| F<sub>2</sub> | 96 | 74 | 22 | 3.4 : 1 | 0.64 |

### 3.2 Double-digest RAD sequencing analysis of the F_2_ population and linkage map development

The ddRAD-seq analysis was used to genotype the F_2_ population for genetic mapping of the locus for the white seed trait derived from ‘ShinanoPower’. The genomic DNA polymorphisms between the two parental lines, ‘ShinanoPower’ and ‘Escort’, were assessed through ddRAD-seq analysis using PacI and NlaIII restriction enzymes (Seki et al., 2020). Illumina HiSeq sequencing of the ddRAD-seq libraries produced 9,281,482 and 8,107,399 single reads (100 bp) for ‘ShinanoPower’ and ‘Escort’ plants, respectively. RAD-tags were extracted from the sequence reads of individual samples. In total, 346,396 and 302,738 RAD-tags with more than two read counts were obtained for the ‘ShinanoPower’ and ‘Escort’ samples, respectively. Comparing the RAD-tags of the two parental lines, 135,129 and 91,471 unique tags were identified as either ‘ShinanoPower’- or ‘Escort’ -specific tags, respectively, whereas 211,267 RAD-tags appeared in both samples. Read mapping was performed with the unique RAD-tags of each parent against the reference lettuce genome sequence [version8 from the crisphead cultivar ‘Salinas’; (https://genomevolution.org/coge/GenomeInfo.pl?gid=28333)] (Reyes-Chin-Wo et al., 2017). A total of 2,871 pairs of RAD-tags (designated as biallelic tags) harboring SNPs or InDels from the two parental lines were described, and these biallelic tags were employed as co-dominant markers for further genetic mapping. The genotypes of these 1,038 biallelic tagged loci were also determined through the ddRAD-seq analysis of 96 individuals in the F_2_ population. Genotypes of the biallelic tag loci in the 96 F_2_ individuals were determined based on the presence or absence of each allelic tag. After excluding loci with missing data, genotyping data from 856 biallelic tag loci of 96 F_2_ individuals were used for linkage map construction (Fig. S2, Table 3). Based on the grouping analysis, the marker loci were distributed into nine linkage groups. Ordering the marker loci in each linkage group resulted in a linkage map comprising 988.8 cM (Fig. S2). Summary statistics for the linkage maps are presented in Table 3. Marker density ranged from 0.5 cM per marker (LG7) to 4.2 cM per marker (LG9). The number of markers ranged from 22 (LG9) to 177 (LG7).

**Table 3.** Summary of integrated lettuce linkage groups.

|  |  | Common tags |  | Escort unique tags |  | ShinanoPower unique tags |  | Linkage construct marker |  | Average interval between markers |
| --- | --- | --- | --- | --- | --- | --- | --- | --- | --- | --- |
| Linkage groups | Total mapped tags | No. RAD-tags | (%) | No. RAD-tags | (%) | No. RAD-tags | (%) | No. bi-allelic tags | Map length (cM) | (cM) |
| LG1 | 37,235 | 15,284 | 41.0 | 8,481 | 22.8 | 13,470 | 36.2 | 138 | 105.5 | 0.8 |
| LG2 | 39,121 | 18,603 | 47.6 | 8,830 | 22.6 | 11,688 | 29.9 | 67 | 43.0 | 0.6 |
| LG3 | 51,405 | 25,399 | 49.4 | 10,965 | 21.3 | 15,041 | 29.3 | 55 | 152.4 | 2.8 |
| LG4 | 73,106 | 36,016 | 49.3 | 14,848 | 20.3 | 22,242 | 30.4 | 113 | 141.0 | 1.2 |
| LG5 | 64,803 | 31,929 | 49.3 | 13,564 | 20.9 | 19,310 | 29.8 | 101 | 177.7 | 1.8 |
| LG6 | 36,432 | 18,005 | 49.4 | 7,143 | 19.6 | 11,284 | 31.0 | 76 | 87.1 | 1.1 |
| LG7 | 36,744 | 17,301 | 47.1 | 7,844 | 21.3 | 11,599 | 31.6 | 177 | 88.6 | 0.5 |
| LG8 | 60,517 | 29,382 | 48.6 | 12,273 | 20.3 | 18,862 | 31.2 | 107 | 100.5 | 0.9 |
| LG9 | 38,504 | 19,348 | 50.2 | 7,523 | 19.5 | 11,633 | 30.2 | 22 | 93.0 | 4.2 |
| Total | 437,867 | 211,267 | 48.2 | 91,471 | 20.9 | 135,129 | 30.9 | 856 | 988.8 | 1.2 |

### 3.3 Fine mapping of the white seed locus and candidate gene analysis

A single locus tightly linked to the white-seed trait was located in LG7 and flanked by two markers (*LG7_v8_48*.*055 Mbp* and *LG7_v8_49*.*864 Mbp*) based on the genotypes of the biallelic RAD-tags. The marker designated as *LG7_v8_49*.*398 Mbp* exhibited complete co-segregation with the white seed trait within the F_2_ population (Fig. 2). Moreover, fine mapping of the target locus was performed using five markers (Table S3) and 84 cultivars (Table S4). Forty-five cultivars had white seeds and the rest had black seeds. Only the *LG7_v8_49*.*251Mbp_HinfI* marker was associated with the white seed phenotype (Tables 4 and S4). A subsequent marker was designed based on a HinfI cut site in the stop codon, *Lsat_1_v5_gn_7_35020*.*1* (Fig. 3). Based on the marker data, it was predicted that the responsible gene was located between 49.173 and 49.326 Mbp in LG7. Eight open reading frames (ORFs) were positioned in this region according to the annotated reference genome sequence of *L. sativa* V8 (Table 5). The sequences of these eight ORFs were compared between ‘ShinanoPower’ and ‘Escort’. There were no small InDels or nonsynonymous substitutions in these seven ORFs between the two parental lines; however, a single nucleotide mutation in a stop codon was found in ORF 7, which is referred to as *Lsat_1_v5_gn_7_35020*.*1*. The allele of the white seed cultivar encoded an additional 78 bp at the 3′end that was not present in the black seed allele (Fig. 3 and Fig. S3). Based on the analysis using publicly available resequencing data from 131 cultivars, there was a complete correlation between stop codon polymorphisms and seed color (Table S5). Phylogenetic analysis revealed that *Lsat_1_v5_gn_7_35020*.*1* is closely related to the *TT2* gene encoding the R2R3-MYB transcription factor (Fig. 4), which is involved in the regulation of seeds color in *Arabidopsis thaliana* (Nesi et al., 2001); we named it as *LsTT2*. R2R3-MYB forms the MYB-bHLH-WDR (MBW) ternary protein complex, together with bHLH-type transcriptional regulators and WD repeat protein (Lepiniec et al., 2006). MBW complex regulates the transcription of gene subsets related to anthocyanin and proanthocyanidin synthesis, thereby modulating the pool size of these metabolites. In addition, the paralog with the greatest similarity, *Lsat_1_v5_gn_5_135961*.*1*, was related to the regulation of anthocyanin biosynthesis in the leaf of red leaf cultivars and was named as Red Lettuce Leaf 2 (RLL2) (Su et al., 2020).

**Table 4.** Seed color and genotype of multiple markers in 84 lettuce cultivars.

| Seed color | Genotype | Expected genotype | Marker name |  |  |  |  |
| --- | --- | --- | --- | --- | --- | --- | --- |
|  |  |  | LG7_v8_48.637 Mbp | LG7_v8_49.173Mbp_EcoRI | LG7_v8_49.251Mbp_Hinfl | LG7_v8_49.326 Mbp | LG7_v8_49.798 Mbp |
| White | ShinanoPower (White seed coat) | 45 | 33 | 34 | 45 | 40 | 41 |
|  | Escort (Black seed coat) | 0 | 12 | 11 | 0 | 5 | 4 |
| Black | ShinanoPower (White seed coat) | 0 | 10 | 7 | 0 | 8 | 4 |
|  | Escort (Black seed coat) | 39 | 29 | 32 | 39 | 31 | 35 |

**Table 5.** Candidate genes in the genomic region between 49.173 Mbp and 49.326 Mbp in LG7.

| ORF | Gene model name | Position of reference genome sequence |  | Putative function | Sequencing |
| --- | --- | --- | --- | --- | --- |
|  |  | Start | End |  |  |
| ORF1 | Lsat_1_v5_gn_7_35140.1 | 49,203,264 | 49,205,806 | Subtilase family protein, putative | Identical |
| ORF2 | Lsat_1_v5_gn_7_35121.1 | 49,225,195 | 49,229,412 | FRIGIDA interacting protein, putative | Identical |
| ORF3 | Lsat_1_v5_gn_7_35101.1 | 49,231,640 | 49,234,256 | Acyl-CoA N-acyltransferases (NAT) superfamily protein, putative | Identical |
| ORF4 | Lsat_1_v5_gn_7_35080.1 | 49,234,299 | 49,235,990 | mitochondrial editing factor, putative | Identical |
| ORF5 | Lsat_1_v5_gn_7_35060.1 | 49,239,808 | 49,242,933 | RNI-like superfamily protein, putative | Identical |
| ORF6 | Lsat_1_v5_gn_7_35041.1 | 49,250,652 | 49,252,131 | TATA-binding related factor (TRF) of subunit 20 of Mediator complex, putative | Identical |
| ORF7 | Lsat_1_v5_gn_7_35020.1 | 49,251,738 | 49,252,982 | R2R3-MYB transcriptional factor, putative | A single nucleotide mutation in stop codon |
| ORF8 | Lsat_1_v5_gn_7_34980.1 | 49,324,772 | 49,325,209 | ELF4-like, putative | Identical |

**Fig. 2.**
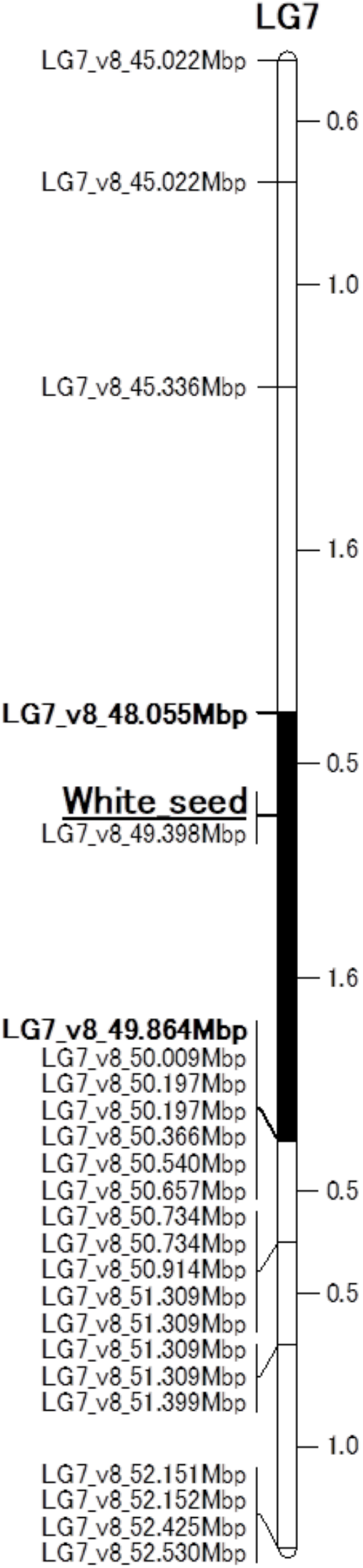
The mapped location of the white seed locus on LG7. Genetic distances (cM) are shown between the markers. “White seed” indicates the position of the responsible gene for the trait of white seed. Black bar indicate white seed locus.

**Fig. 3.**
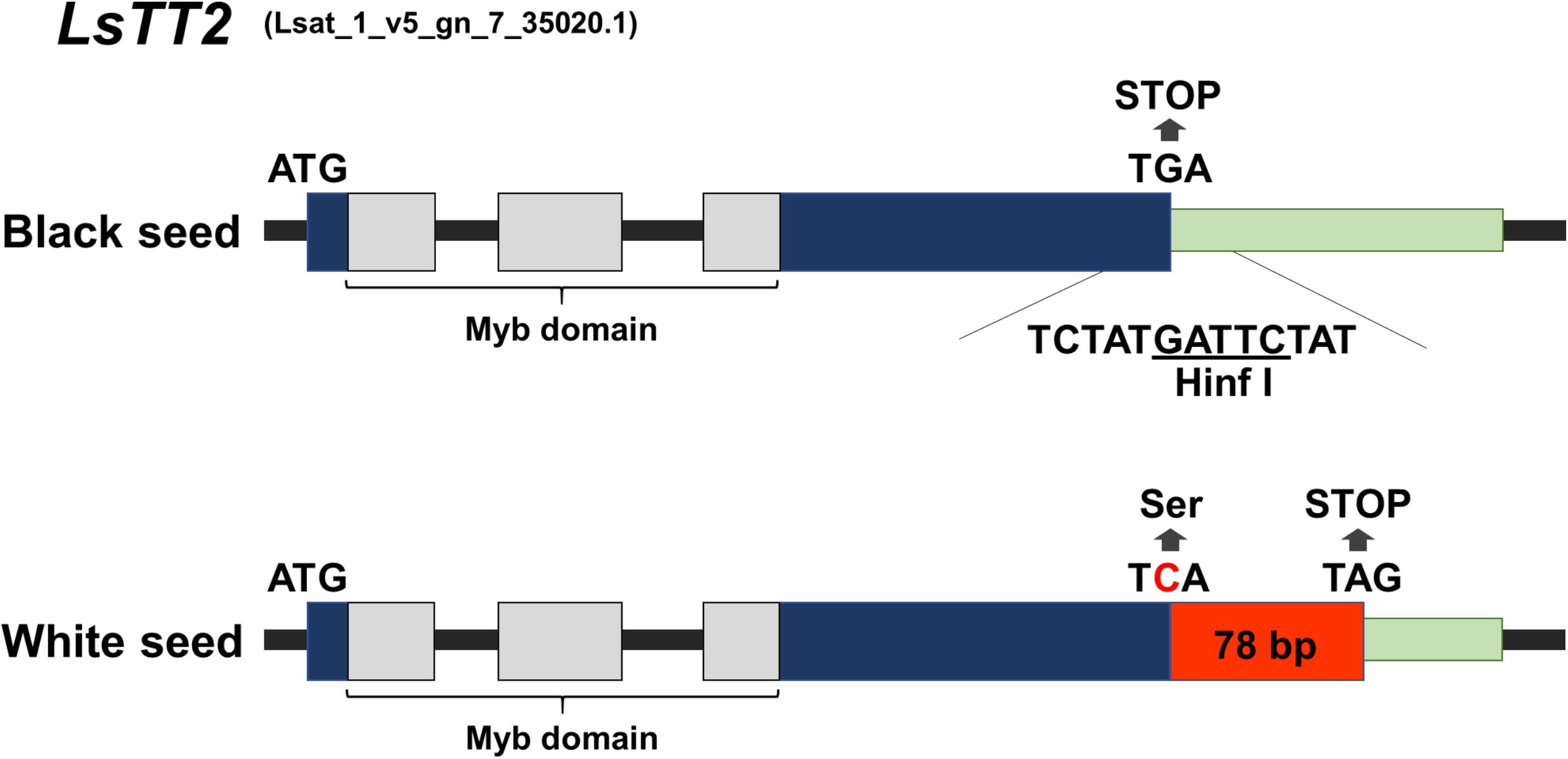
Comparison of *Lsat_1_v5_gn_7_35020*.*1* between black and white seeds. This white seed allele encodes an additional 78 bp at the 3′ end that are not present in black seed allele. The black seed allele sequence contains a Hinf I restriction site in the stop codon.

**Fig. 4.**
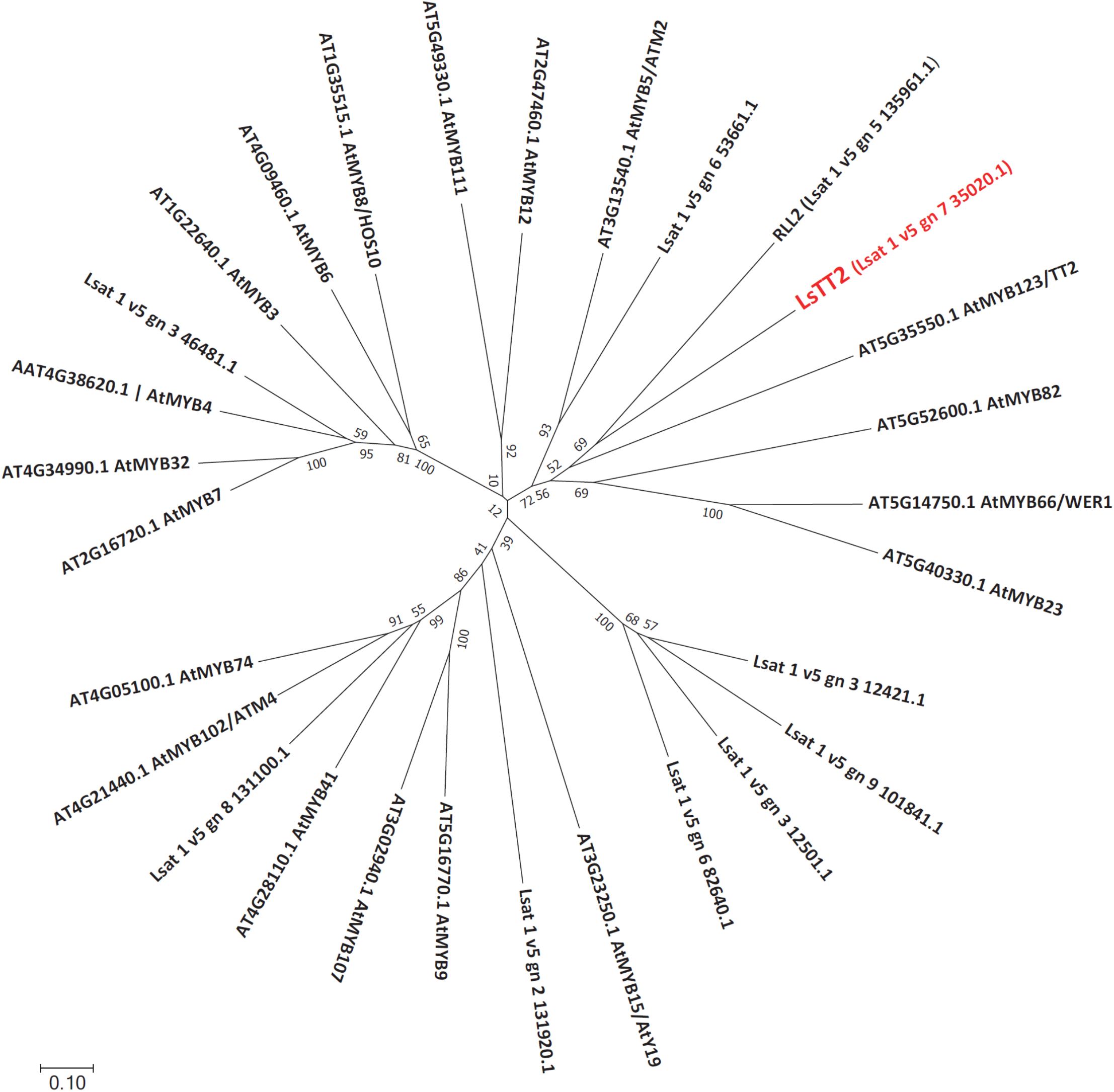
Phylogenetic analysis of the MYB domain-containing proteins in *L. sativa*. The evolutionary tree was built with orthologs encoding MYB transcription factor in lettuce and *Arabidopsis*. The red font indicates the candidate gene controlling seed color in lettuce.

### 3.4 Analysis of pigment accumulation in seeds

The absorbances of the supernatants extracted from black and white seeds at 500 nm were 0.061 (SE = 0.015) and 0.038 (SE = 0.008), respectively. The proanthocyanidin content in the white seed was 0.62 times lower than that in the black seeds. Some phenylpropanoids such as chlorogenic acid were found in both seed colors; however, common anthocyanins and other flavonoids were not detected in the HPLC analysis (Fig. S4 and S5). White seeds appear to have reduced function with regard to the accumulation of proanthocyanidins.

### 3.5 Validation of *LsTT2* function through CRSPR/Cas9-based genome editing

To obtain knockout strains with a loss-of-function mutation in the *LsTT2* gene, gRNA was designed at the first exon for generating the early stop codon. CRISPR/Cas9 vector was introduced via Agrobacterium into the lettuce, ‘Oilseed’, which normally produces black seeds. The transformed plants were selected for both antibiotic resistance and PCR-positive results for foreign genes. Nine acclimated T_0_ individuals were used for the analysis of the target sequence. The resulting sequence variations of the eight clones allowed us to predict the genotype of the strains: monoallelic or biallelic, heterozygous or homozygous, in-frame or out-of-frame. Six strains produced white seeds in the T_0_ phenotype (Table 6). Among these, three strains (4-1, 15-1, 19-2) could be biallelic homozygous mutants with a single-base insertion. The other three strains (5-1, 30-4, 22-2) that produced white seeds were biallelic heterozygous mutants with insertions of one or two bases. All edits generated early stop codons and 13 (MGRSPCLFKDWSE*) or 12 (MGRSPCLVQRLV*) short peptides. The three remaining strains (15-2, 25-4, 19-1) had black seeds, despite genome editing. This could be attributed to the presence of biallelic heterozygous mutants, including in-frame editing with a three-base deletion. These results confirmed that *LsTT2* controls the dominant traits of seed color. The inner parts when observed without the pericarp were brown in all genotypes (Fig. 5), suggesting that *LsTT2* is involved only in achene color.

**Table 6.**
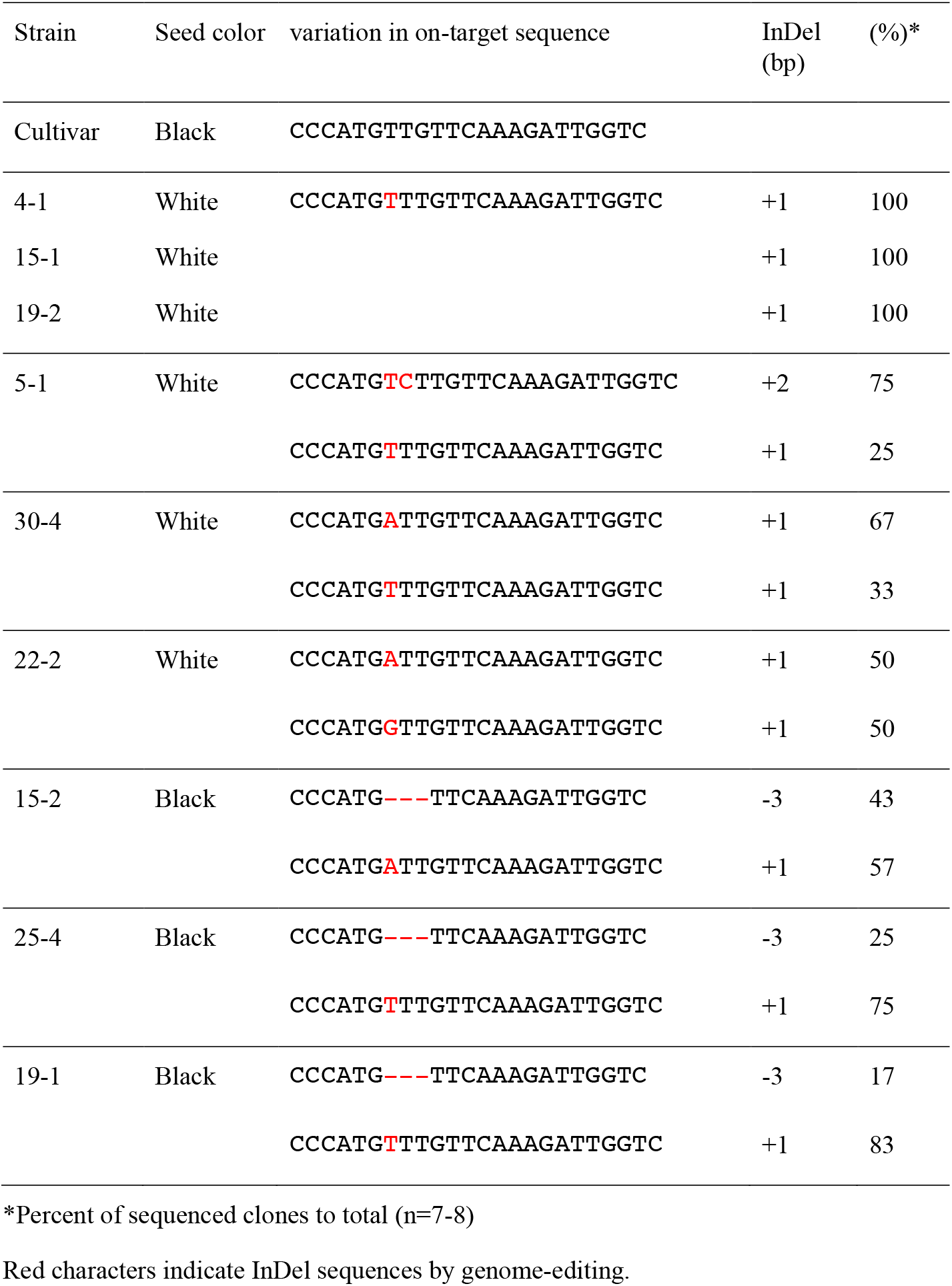
Seed color and sequence variation of genome-edited *LsTT2* gene in T_0_ plants.

**Fig. 5.**
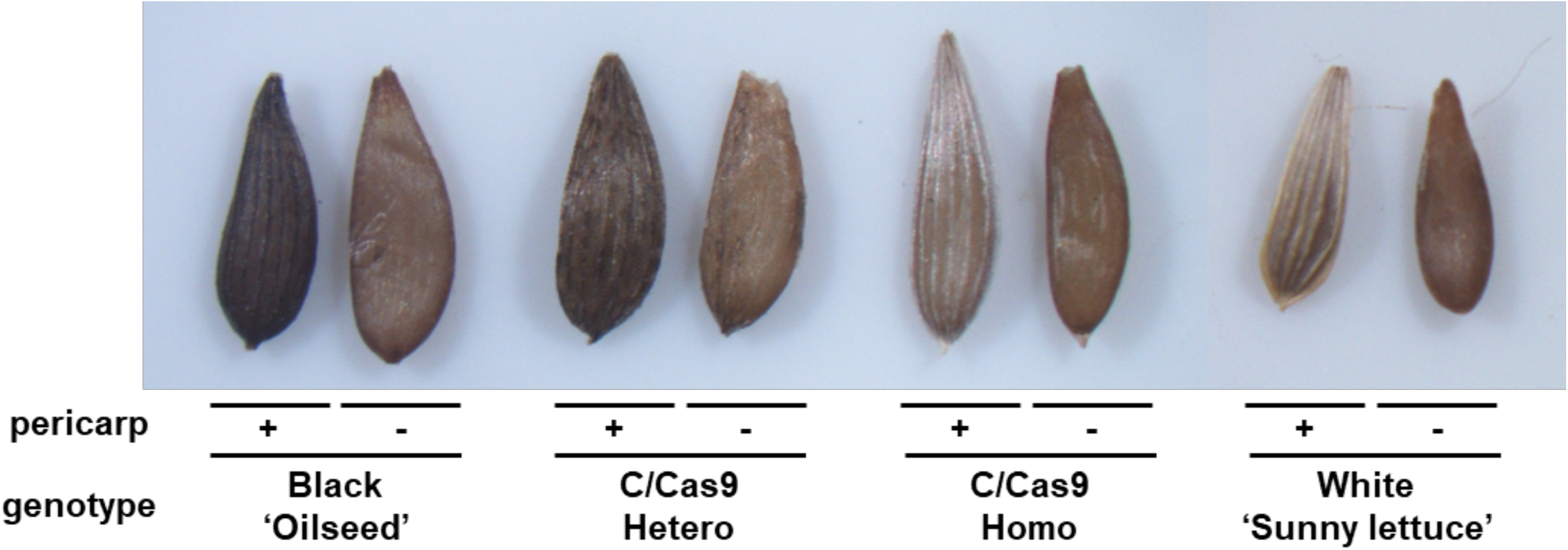
Characterization of seed color of genome-edited lettuce. The outermost appearance was determined alternatively as black or white. Pericarp was removed from lettuce achene (-) or not (+). ‘Oilseeds’ is a black seed cultivar, and T_1_ seed is from genome-edited plants (C/Cas9), while ‘Sunny lettuce’ is a white seed cultivar. The seed color described here is derived from the parental T_0_ trait.

## 4 Discussion

In this study, we succeeded in the genetic mapping and identification of the genes responsible for white seeds in lettuce. According to genetic mapping using ddRAD-seq and PCR-based markers, the locus of white seeds was located between 49.173 Mbp and 49.326 Mbp in LG7 (Fig. 2 and Table 4). Eight predicted genes were identified in this region (Table 5). Resequence analysis between ‘ShinanoPower’ (white seed) and ‘Escort’ (black seed) revealed that the candidate genes of the coding sequences were identical except for *LsTT2*, gene model name *Lsat_1_v5_gn_7_35020*.*1* (Table 5). *LsTT2* has a single nucleotide mutation in the stop codon of cultivars with white seeds, and the *LG7_v8_49*.*251Mbp_HinfI* marker, which employs this mutation, was completely linked to the white seed phenotype (Fig. 3 and Table 4). Analysis using publicly available resequencing data showed a complete correlation between stop codon polymorphisms and seed color (Table S4). In addition, this marker position overlaps with a previously reported locus for white (w) seed color (Kwon et al., 2013; Simko et al., 2013). Pigment accumulation in plants is controlled by two gene subsets: early biosynthetic genes and late biosynthetic genes (LBGs) (Kubasek et al., 1992; Quattrocchio et al., 1993; Nesi et al., 2000). The transcription factor complex, consisting of *AtTT2, AtTT8*, and *AtTTG1* controls the expression of LBGs (Gonzalez et al., 2008, 2016), and *AtTT2* mutants have altered seed color into yellow due to no proanthocyanidin production in *Arabidopsis* (Shirley et al., 1995; Nesi et al., 2001). AtTT2 controls the expression of *BAN* (anthocyanidin reductase gene) involved in the divergence of proanthocyanidins and anthocyanins during flavonoid biosynthesis (Debeaujon et al., 2003). Therefore, as in Arabidopsis, proanthocyanidins rather than anthocyanins may be responsible for seed color in lettuce (Fig. S4). LsTT2 shared 28% identity with AtTT2, and highly conserved DNA-binding domain containing R2 and R3 repeats, which consists of ‘-W-(X19)-W-(X19)-W-’ and ‘-F/I-(X18)-W-(X18)-W-’, respectively. Therefore, *LsTT2* is a biologically plausible candidate gene. Genome editing of the target, *LsTT2* showed that knockout mutants harboring an early termination codon produced white seeds (Table 6, Fig. 5). We infer that the orthologous proteins in lettuce probably have a conserved function; the mutation of the stop codon of *LsTT2* in white seeds causes a significant conformational change and interferes with complex formation with other interactions. Dominant-negative effects of MYB due to deletions in the C-terminal end have been reported in Arabidopsis (Velten et al., 2010). Within the amino acid sequence of an additional 26 residues at the C-terminus, we could not find any typical repressor motifs for MYB, such as the EAR (ERF-associated amphiphilic repression; LxLxL or DLNxxP), SID (Sensitive to ABA and Drought 2 protein interact motif; GY/FDFLGL), or TLLLFR (Wu et al., 2022). It is worth exploring the analysis of additional sequences, including the possibility of identifying novel inhibitory motifs. The white seeds exhibited reduced accumulation of proanthocyanidins compared to the black seeds based on the vanillin–sulfuric acid assay, which is in agreement with the conclusion that the white seed cultivars have a reduced-function mutation in *LsTT2* that controls the expression of LBGs in lettuce seeds. In conclusion, *LsTT2* is the allele responsible for the shift in seed color from black to white.

During the artificial crossing of lettuce flowers, the breeder must remove the maternal parent pollen from the flower. However, it is impossible to completely remove pollen from lettuce flowers; therefore, seeds of both the selfed progeny and the F_1_ hybrid are produced unintentionally owing to a compound autogamous floral structure (Simko et al., 2011). Therefore, breeders would like to utilize the inheritance patterns of easily recognizable traits to distinguish F_1_ hybrids from selfed plants in the following progeny. Following a cross between white the seed pure line (♀) and black seed pure line (♂), selfed progenies produce the next generation of white seed and F_1_ hybrids produce the F_2_ generation of black seed because of the dominant trait (Thompson, 1942; Ryder, 1999). Therefore, seed color can be effectively used to distinguish between selfed and hybrid plants (Thompson, 1942). The marker used to distinguish between F_1_ hybrids and selfed plants in populations derived from the white seed × black seed cross could also contribute to validating the phenotype of the F_1_ generation. The verification of inconspicuous traits that require bioassays, such as disease resistance, has been difficult in the F_1_ generation. Male sterility makes it possible to produce only F_1_ hybrid seeds (Hayashi et al., 2011; Seki, 2022); however, it has not been widely used because of limited cross combinations. The inheritable characteristics of the F_1_ generation can be examined using a bioassay with only the seeds of the hetero genotype. From the F_2_ generation onward, it was possible to intentionally select the seed color using the marker. These approaches are valuable for the development of breeding methods that accelerate the development of lettuce cultivars. Therefore, *LG7_v8_49*.*251Mbp_HinfI* marker-targeted *LsTT2* could be used to distinguish almost all white seeds of lettuce worldwide and could be applied to significantly enhance lettuce breeding programs (Tables S1, S2, and S3).

Lettuce was first domesticated near the Caucasus because of the loss of seed-shattering by spontaneous mutation (Wei et al., 2021). Therefore, cultivated lettuce has a non-seed-shattering due to the same *qSHT* locus. Because the domestication time is estimated to be around 4000 _BC_, the change in seed color is plus onwards. Considering the fact that lettuce was depicted on wall paintings of Egyptian tombs around 2500 _BC_ as one of the major vegetable crops (De Vries, 1997), it is reasonable to assume that the change in seed color occurred before the spread of lettuce seed around the world (Tables S4 and S5). The discovery of the white seed, which has been used as an important agronomic trait for thousands of years, is indeed a great achievement.

## 5 Conclusion

The development of a robust marker for marker-assisted selection and identification of the gene responsible for white seeds has implications for the lettuce breeding and agricultural aspects of seed color. This study not only identified a gene responsible for the white seed, but also revealed an important gene for a key domestication trait for lettuce cultivation and breeding. These findings could be useful for future lettuce-breeding endeavors.

## Conflict of Interest

The authors declare that they have no conflict of interests.

## 7 Author Contributions

KS, KK, and YU designed experiments. KS developed the mapping population and the ddRAD-seq library. KS and KK performed the sequence analysis. KS and YU performed genotyping. KS, KK, and YM collected the seed observation data. YM performed pigment analysis. KY, YU, RK, and KN performed the genome editing. KS, KK, and YU performed data analysis. KS, KK, YM and YU prepared the manuscript. All the authors have read and approved the final version of the manuscript.

## 8 Funding

This study was supported by grants from the Ministry of Agriculture, Forestry and Fisheries of Japan (Project for Climate Change, Vegetable-4103) to KS.

## 9 Acknowledgments

The authors would like to thank Yoko Takahashi, Yoshie Nakayama, and especially Hideaki Okazawa for their technical assistance in the field experiments.

## 10 Data Availability Statement

Raw sequence data (FASTQ) for the ddRAD-seq dataset were deposited in the DNA Data Bank of Japan (DDBJ) Sequence Read Archive (http://ddbj.nig.ac.jp/dra/index_e.html) under the accession number DRA013652.

## Notes

http://ddbj.nig.ac.jp/dra/index_e.html

### Competing Interest Statement

The authors have declared no competing interest.

## References

Borthwick, H. A., Hendricks, S. B., Parker, M. W., Toole, E. H., and Toole, Vi. K. (1952). A Reversible photoreaction controlling seed germination. Proc. Natl. Acad. Sci. 38, 662–666. doi: 10.1073/pnas.38.8.662.

Debeaujon, I., Nesi, N., Perez, P., Devic, M., Grandjean, O., Caboche, M., and Lepiniec, L. (2003). Proanthocyanidin-accumulating cells in Arabidopsis testa: regulation of differentiation and role in seed development. Plant Cell 15, 2514–2531. doi: 10.1105/tpc.014043.

De Vries, I. M. (1997). Origin and domestication of Lactuca sativa L. Genet. Resour. Crop Evol. 44, 165–174. doi: 10.1023/A:1008611200727.

Gonzalez, A., Brown, M., Hatlestad, G., Akhavan, N., Smith, T., Hembd, A., et al. (2016). TTG2 controls the developmental regulation of seed coat tannins in Arabidopsis by regulating vacuolar transport steps in the proanthocyanidin pathway. Dev. Biol. 419, 54–63. doi: 10.1016/j.ydbio.2016.03.031.

Gonzalez, A., Zhao, M., Leavitt, J. M., and Lloyd, A. M. (2008). Regulation of the anthocyanin biosynthetic pathway by the TTG1/bHLH/Myb transcriptional complex in Arabidopsis seedlings. Plant J. 53, 814–827. doi: 10.1111/j.1365-313X.2007.03373.x.

Hayashi, M., Ujiie, A., Serizawa, H., Sassa, H., Kakui, H., Oda, T., et al. (2011). Development of SCAR and CAPS markers linked to a recessive male sterility gene in lettuce (Lactuca sativa L.). Euphytica 180, 429–436. doi: 10.1007/s10681-011-0417-y.

Lepiniec, L., Debeaujon, I., Routaboul, J. M., Baudry, A., Pourcel, L., Nesi, N., Caboche, M. (2006) Genetics and biochemistry of seed flavonoids. Annual Review of Plant Biology 57, 405–430.

Kubasek, W. L., Shirley, B. W., McKillop, A., Goodman, H. M., Briggs, W., and Ausubel, F. M. (1992). Regulation of flavonoid biosynthetic genes in germinating Arabidopsis seedlings. Plant Cell 4, 1229–1236. doi: 10.2307/3869409.

Kwon, S., Simko, I., Hellier, B., Mou, B., and Hu, J. (2013). Genome-wide association of 10 horticultural traits with expressed sequence tag-derived SNP markers in a collection of lettuce lines. Crop J. 1, 25–33. doi: 10.1016/j.cj.2013.07.014.

Li, H., and Durbin, R. (2009). Fast and accurate short read alignment with Burrows-Wheeler transform. Bioinformatics 25, 1754–1760. doi: 10.1093/bioinformatics/btp324.

Nesi, N., Debeaujon, I., Jond, C., Pelletier, G., Caboche, M., and Lepiniec, L. (2000). The TT8 gene encodes a basic helix-loop-helix domain protein required for expression of DFR and BAN genes in Arabidopsis siliques. Plant Cell 12, 1863–1878. doi: 10.1105/tpc.12.10.1863.

Nesi, N., Jond, C., Debeaujon, I., Caboche, M., and Lepiniec, L. (2001). The Arabidopsis TT2 gene encodes an R2R3 MYB domain protein that acts as a key determinant for proanthocyanidin accumulation in developing seed. Plant Cell 13, 2099–2114. doi: 10.1105/tpc.13.9.2099.

Quattrocchio, F., Wing, J. F., Leppen, H. T. C., Mol, J. N. M., and Koes, R. E. (1993). Regulatory genes controlling anthocyanin pigmentation are functionally conserved among plant species and have distinct sets of target genes. Plant Cell 5, 1497–1512. doi: 10.1105/tpc.5.11.1497.

Reyes-Chin-Wo, S., Wang, Z., Yang, X., Kozik, A., Arikit, S., Song, C., et al. (2017). Genome assembly with in vitro proximity ligation data and whole-genome triplication in lettuce. Nat. Commun. 8, 1–11. doi: 10.1038/ncomms14953.

Robinson, J., Thorvaldsdóttir, H., and Winckler, W. (2011). Integrative genomics viewer. Nat Biotechnol 29, 24–26.

Ryder, E. J. (1999). Lettuce, Endive and Chicory. CABI Publishing, Wallingford.

Seki, K. (2021). RAD-R scripts: R pipeline for RAD-seq from FASTQ files to linkage maps construction and run R/QTL, operating only at copying and pasting scripts into R console. Breed. Sci. doi: 10.1270/jsbbs.20159.

Seki, K. (2022). Detection of candidate gene LsACOS5 and development of InDel marker for male sterility by ddRAD-seq and resequencing analysis in lettuce. Sci. Rep. 12, 1–8. doi: 10.1038/s41598-022-11244-2.

Seki, K., Komatsu, K., Tanaka, K., Hiraga, M., Kajiya-Kanegae, H., Matsumura, H., et al. (2020). A CIN-like TCP transcription factor (LsTCP4) having retrotransposon insertion associates with a shift from Salinas type to Empire type in crisphead lettuce (Lactuca sativa L.). Hortic. Res. 7, 1–14. doi: 10.1038/s41438-020-0241-4.

Shimatani, Z., Kashojiya, S., Takayama, M. et al. (2017). Targeted base editing in rice and tomato using a CRISPR-Cas9 cytidine deaminase fusion. Nat. Biotechnol. 35, 441–443. 10.1038/nbt.3833

Shirley, B. W., Kubasek, W. L., Storz, G., Bruggemann, E., Koornneef, M., Ausubel, F. M., et al. (1995). Analysis of Arabidopsis mutants deficient in flavonoid biosynthesis. plant J. 8, 659–671.

Simko, I., Atallah, A. J., Ochoa, O. E., Antonise, R., Galeano, C. H., Truco, M. J., et al. (2013). Identification of QTLs conferring resistance to downy mildew in legacy cultivars of lettuce. Sci. Rep. 3, 1–10. doi: 10.1038/srep02875.

Simko, I., Hayes, R. J., Truco, M. J., and Michelmore, R. W. (2011). Mapping a dominant negative mutation for triforine sensitivity in lettuce and its use as a selectable marker for detecting hybrids. Euphytica 182, 157–166. doi: 10.1007/s10681-011-0407-0.

Su, W., Tao, R., Liu, W., Yu, C., Yue, Z., He, S., et al. (2020). Characterization of four polymorphic genes controlling red leaf colour in lettuce that have undergone disruptive selection since domestication. Plant Biotechnol. J. 18, 479–490. doi: 10.1111/pbi.13213.

Thompson, R. C. (1942). Inheritance of seed color in lactuca sativa. J. Agric. Res. 66, 441–446.

Velten, J., Cakir, C., and Cazzonelli, C. I. (2010) A spontaneous dominant-negative mutation within a 35S::AtMYB90 transgene inhibits flower pigment production in tobacco. PLoS ONE 5, e9917. doi:10.1371/journal.pone.0009917

Voorrips, R. E. (2002). MapChart: Software for the graphical presentation of linkage maps and QTLs. J. Hered. 93, 77–78. doi: 10.1093/jhered/93.1.77.

Wang, Y., Lu, H., and Hu, J. (2016). Molecular mapping of high resistance to bacterial leaf spot in lettuce PI 358001-1. Phytopathology 106, 1319–1325. doi: 10.1094/PHYTO-09-15-0238-R.

Waycott, W., Fort, S. B., Ryder, E. J., and Michelmore, R. W. (1999). Mapping morphological genes relative to molecular markers in lettuce (Lactuca sativa L.). Heredity (Edinb). 82, 245–251. doi: 10.1038/sj.hdy.6884730.

Wei, T., van Treuren, R., Liu, X., Zhang, Z., Chen, J., Liu, Y., et al. (2021). Whole-genome resequencing of 445 Lactuca accessions reveals the domestication history of cultivated lettuce. Nat. Genet. 53, 752–760. doi: 10.1038/s41588-021-00831-0.

Widell, K.O. and Vogelmann, T.C. (1988). Fiber optic studies of light gradients and spectral regime within Lactuca sativa achenes. Physiol. Plantaarum 72, 706–712.

Woolley, J. T., and Stoller, E. W. (1978). Light penetration and light-induced seed germination in soil. Available at: http://www.plantphysiol.org.

Wu,Y., Wen, J., Xia, Y., Zhang, L., and Du, H. (2022) Evolution and functional diversification of R2R3-MYB transcription factors in plants, Hort. Res. 9, uhac058, 10.1093/hr/uhac058

