## Supplemental Tables and Figures for "*LsTT2* encoding R2R3-MYB transcription factor is responsible for a shift from black to white in lettuce seed"

### 1 Supplementary Figures

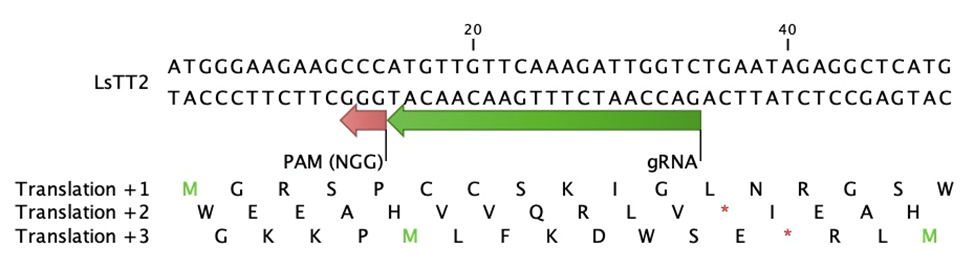

Additional file 1: Fig. S1. *LsTT2* nucleotide sequence and gRNA design. The 50 bp of the first exon of *LsTT2* is shown. The red arrow indicates the PAM sequence and the green arrow indicates gRNA. Indels other than multiples of three cause frameshifts and the appearance of an early termination codon.

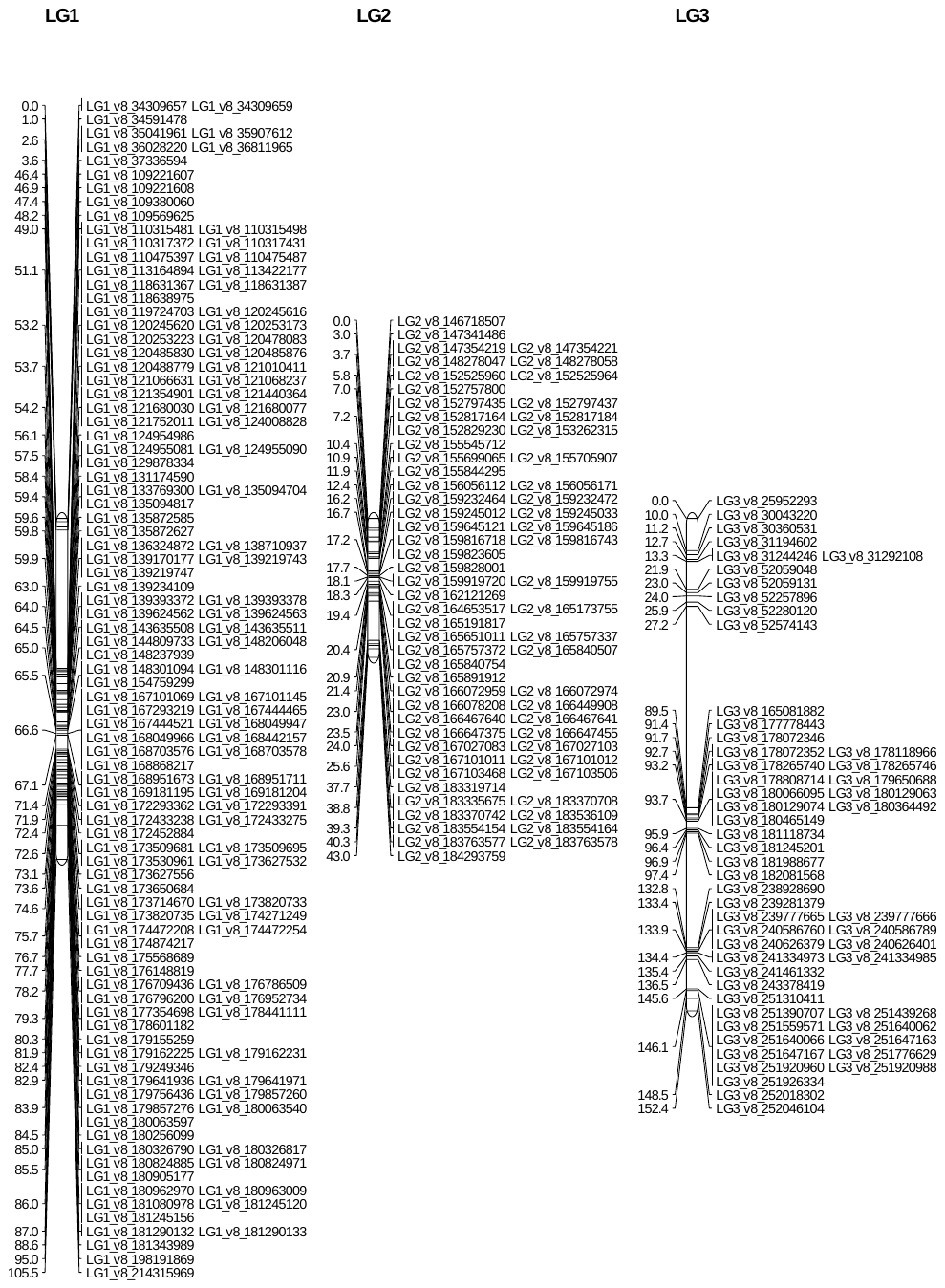

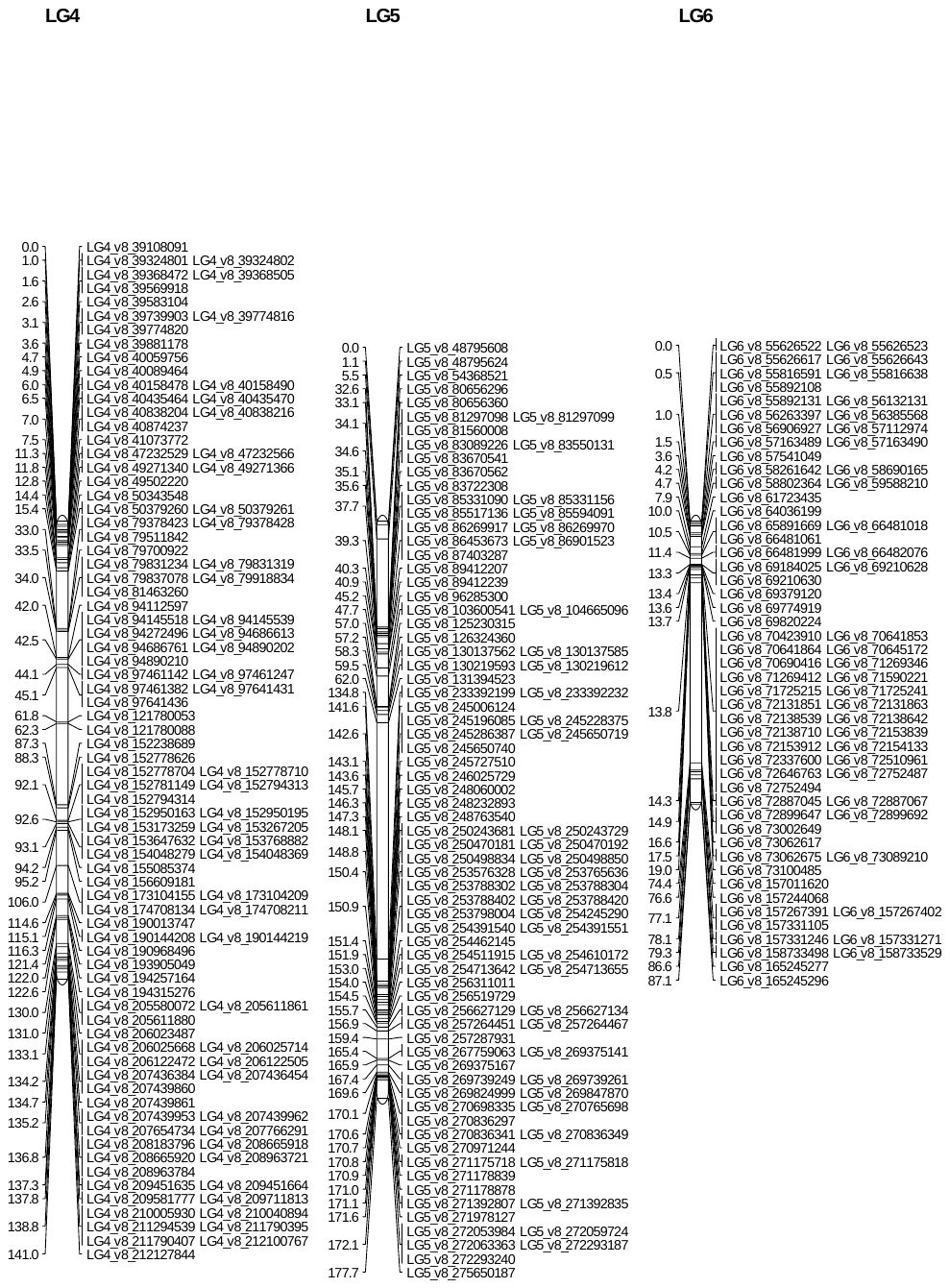

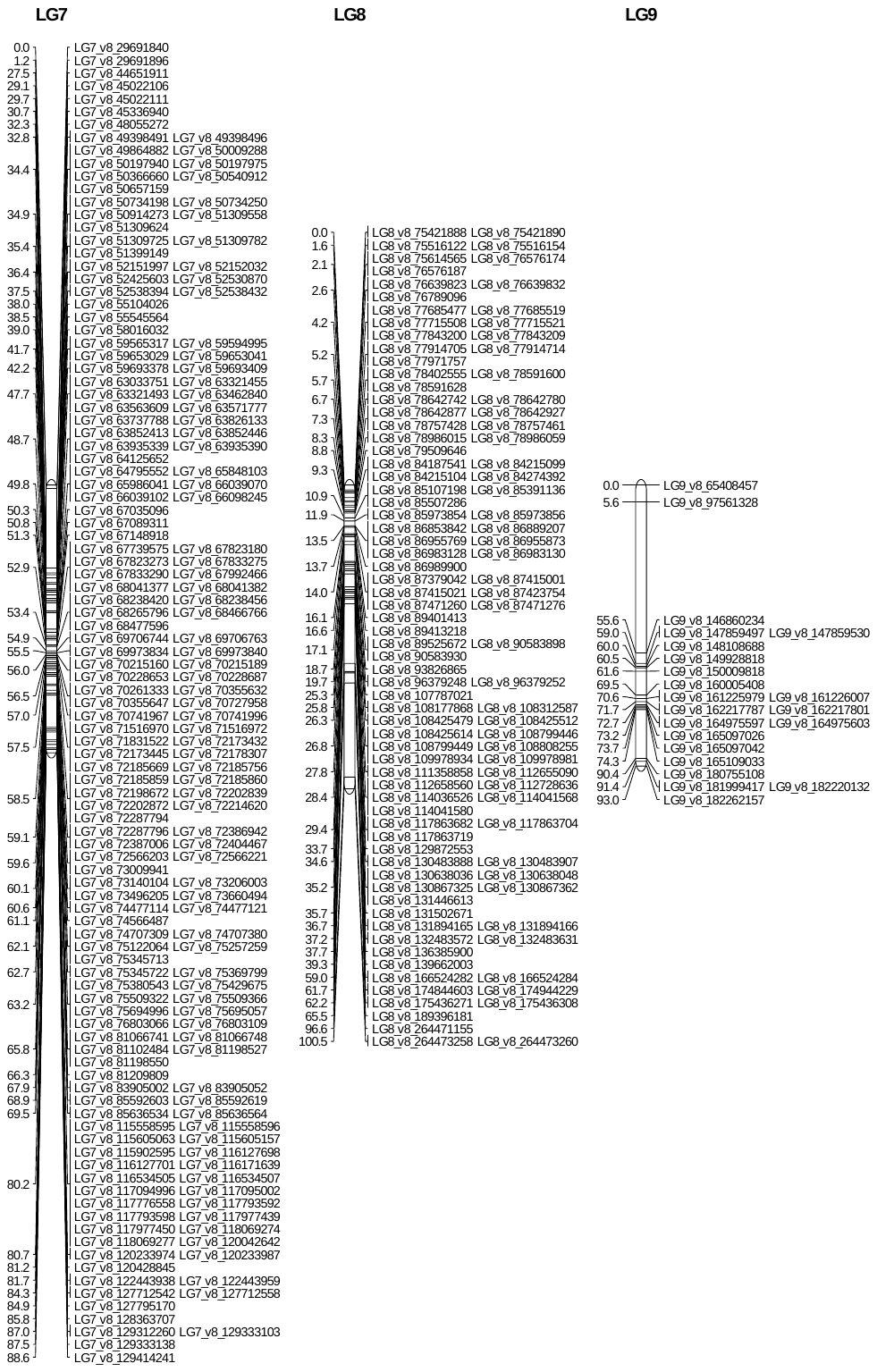

Additional file 2: Fig. S2. Schematic of the consensus map for an F_2_ population derived from a cross between ‘ShinanoPower’ and ‘Escort’. The number on the left indicates the distance in cM and the characters on the right indicate the marker name. Horizontal lines across chromosomes indicate the positions of the loci on each chromosome.

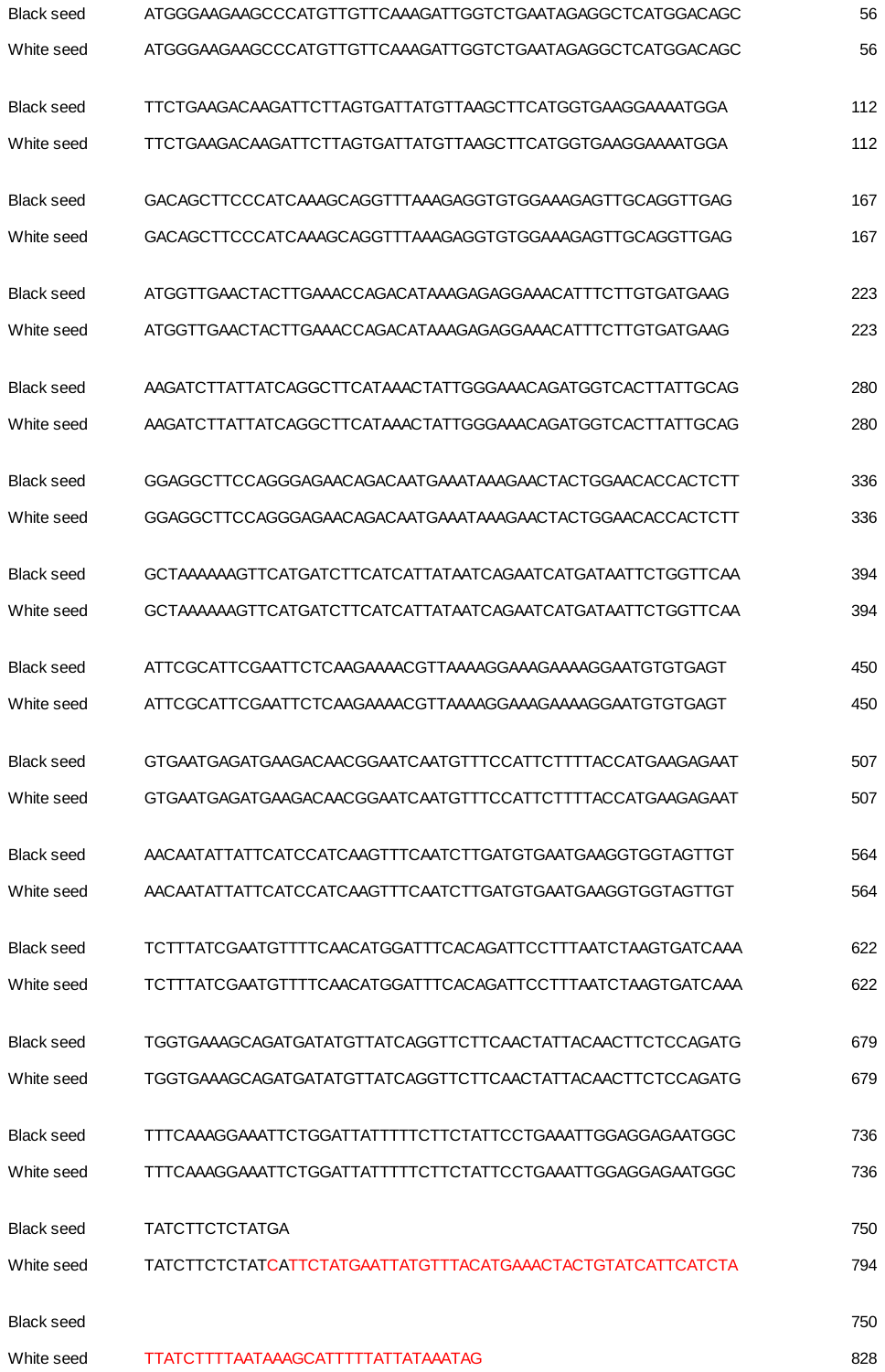

Additional file 3: Fig. S3. Comparison of *LsTT2* nucleic acid sequences of black and white seeds.

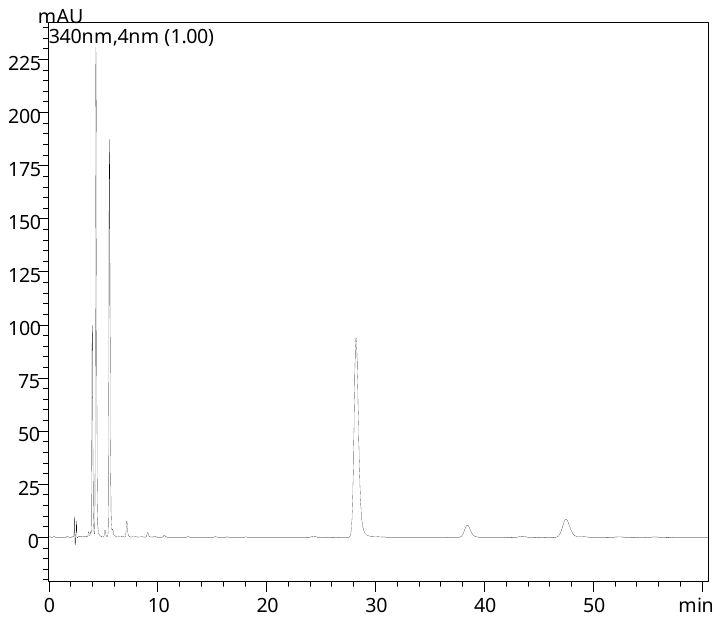

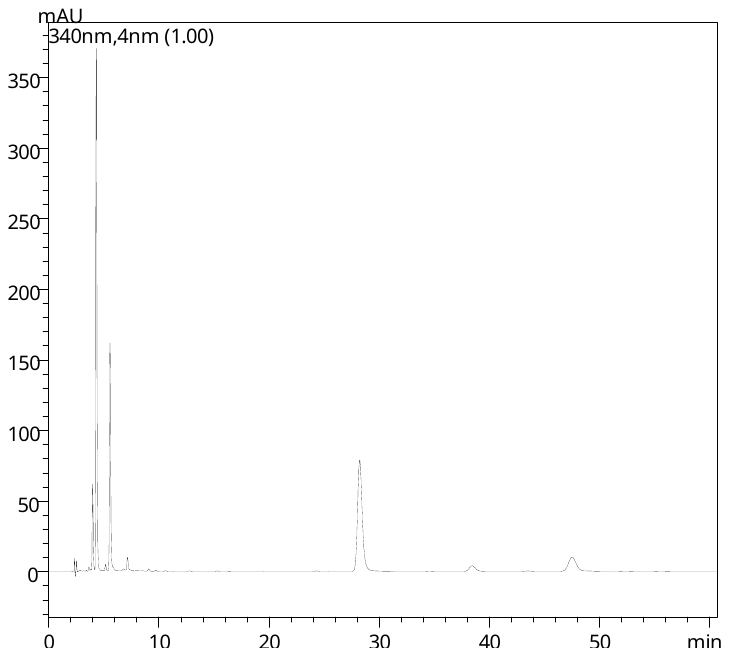

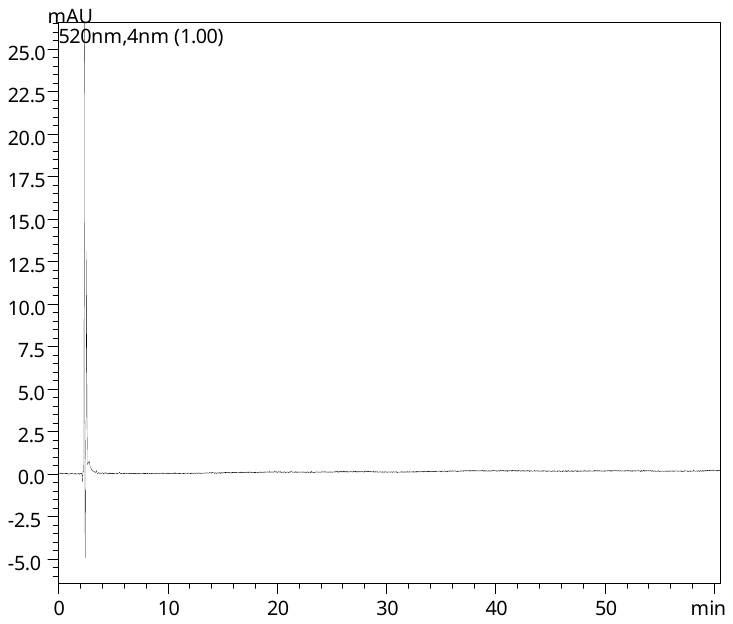

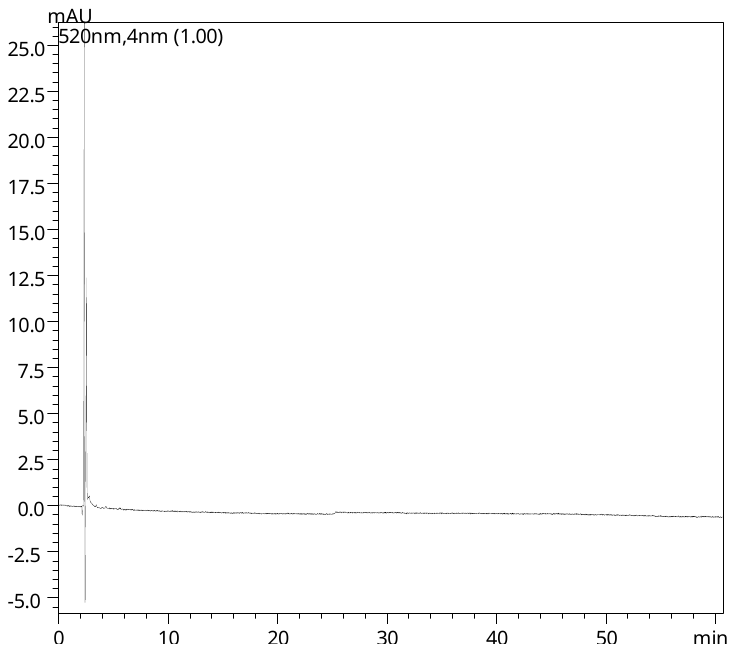

Chlorogenic acid

Chlorogenic acid

White seed (340 nm)

Black seed (340 nm)

White seed (520 nm)

Black seed (520 nm)

Additional file 4: Fig. S4. HPLC chromatograms of black and white seed extracts. Detection of 520 nm was for common anthocyanins, and detection of 340 nm was for other flavonoids and phenylpropanoids. Injection peaks: 2.4-2.8 min, chlorogenic acid: 5.6 min, and other phenylpropanoids: 4.0, 4.3, 7.2, 28.3, 38.5 and 47.6 min.

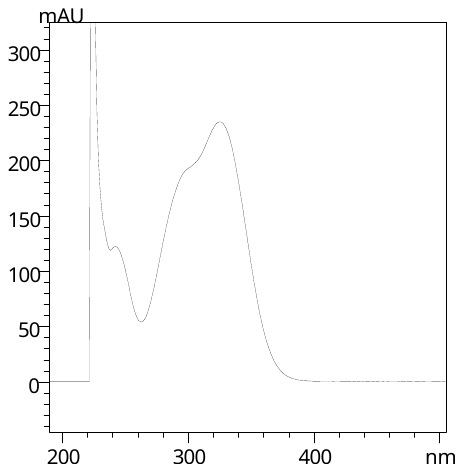

Additional file 5: Fig. S5. UV-vis spectrum of chlorogenic acid (5.6 min) detected from lettuce seed in HPLC survey of this study. The spectra of the other phenylpropanoids were similar.

Additional file 6: Table S1. Number of accessions of black and white seed cultivars organized by country preserved in CGN^*1^ and GRIN^*2^.

^*1^CGN: The Centre for Genetic Resource, the Netherlands

^*2^GRIN: The Germplasm Resources Information Network managed by USDA

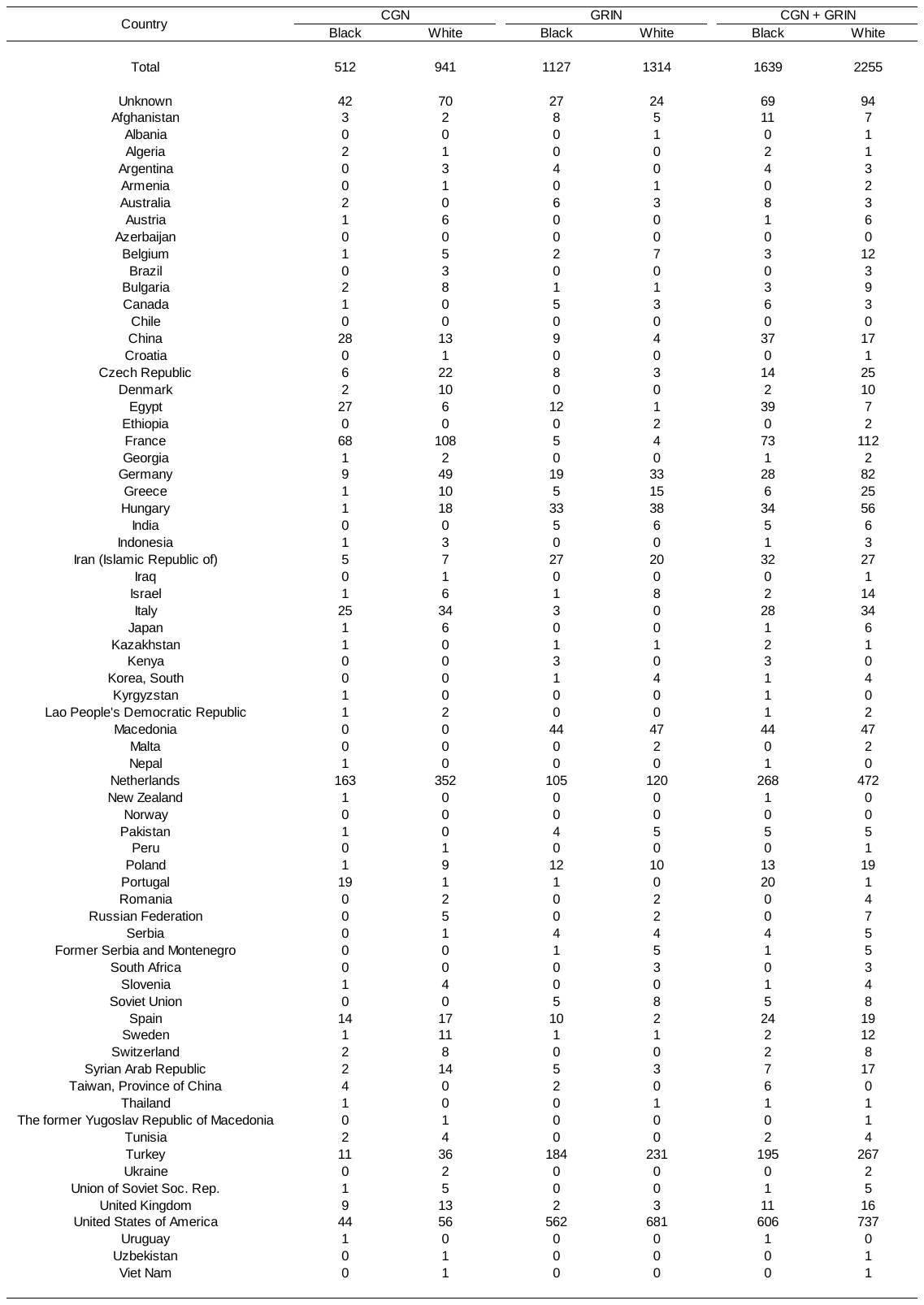

Additional file 7: Table S2. Primer sequences for gRNA.

| *Primer name* | Sequence |
| --- | --- |
| *LsTT2* gRNA primer-F | 5’- ATTGGACCAATCTTTGAACAACAT -3’ |
| *LsTT2* gRNA primer-R | 5’- AAACATGTTGTTCAAAGATTGGTC -3’ |
| Blue letters indicate the protruding ends of the restriction enzyme BbsⅠ. | |

Additional file 8: Table S3. Markers developed within LG7 based on PCR for distinguishing ‘ShinanoPower’ from ‘Escort’.

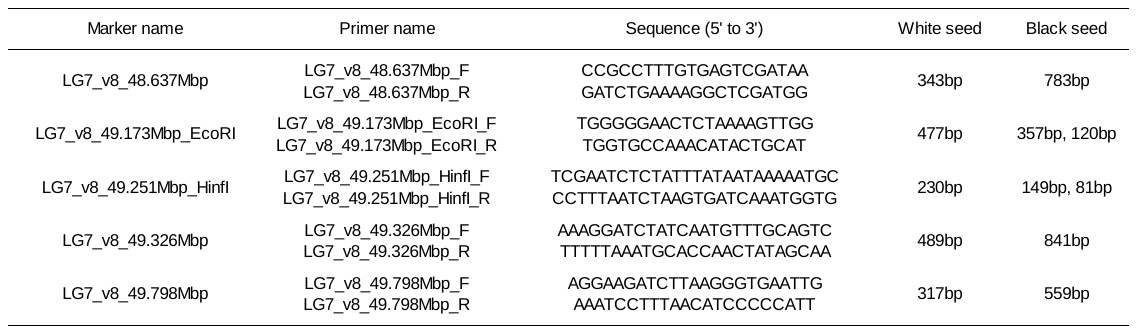

Additional file 9: Table S4. Seed color phenotypic observations for 84 cultivars compared with the predicted colors derived from the genotypic information of the five markers.

Details of the phenotypes and their respective markers are described in Table S3.

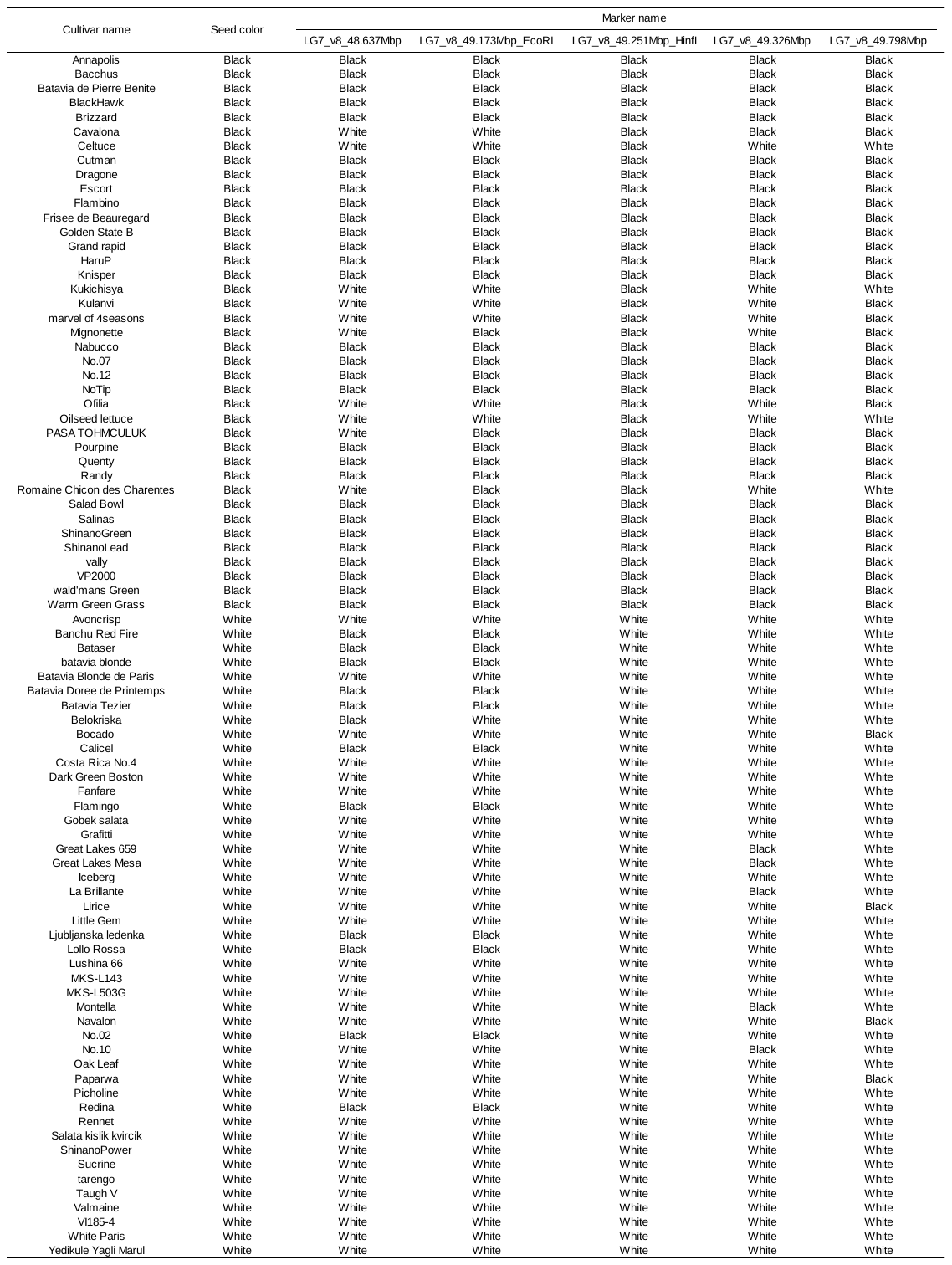

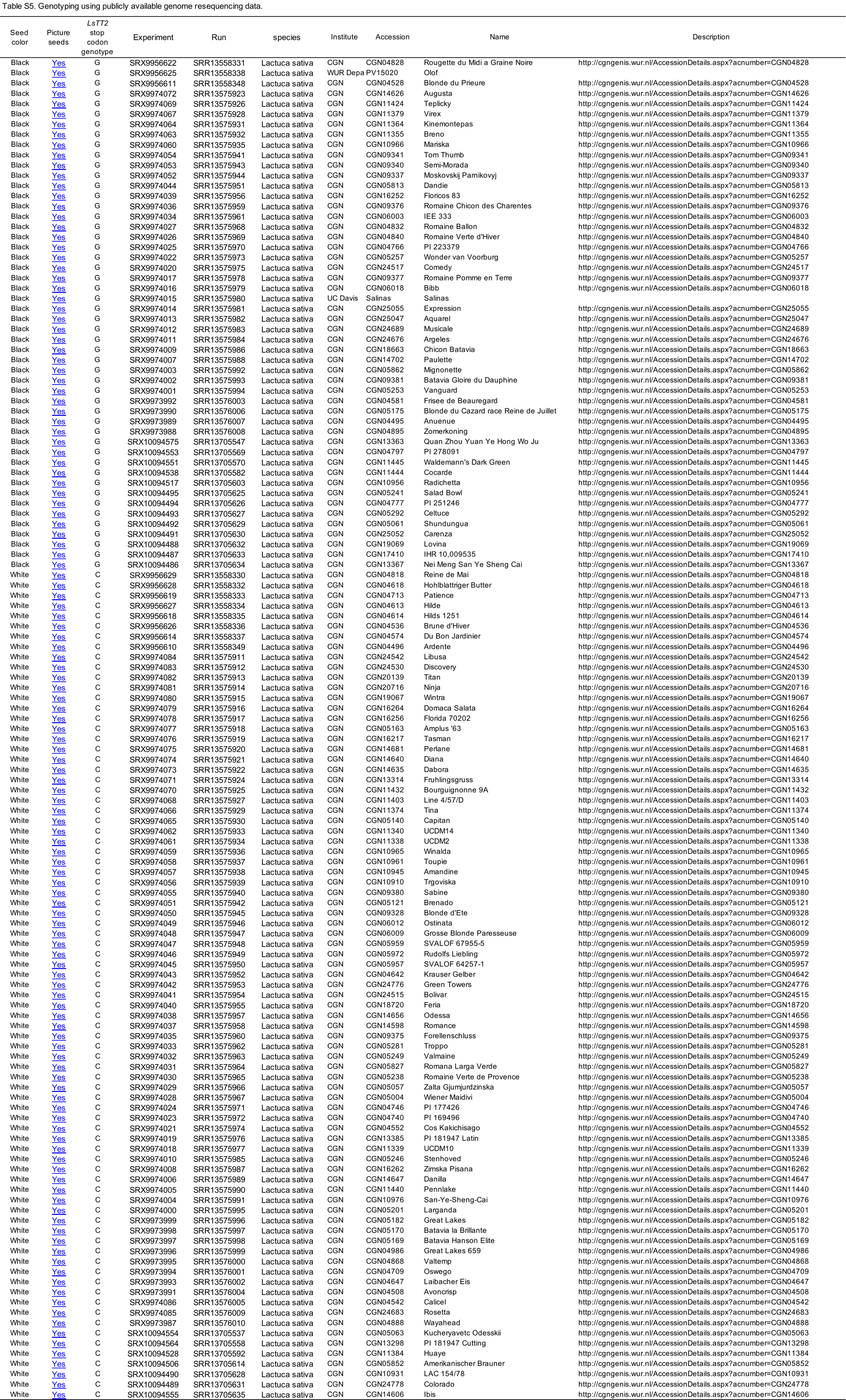
Additional file 10: Table S5. Genotyping using publicly available genome resequencing data.
